# The case against full probability distributions in perceptual decision making

**DOI:** 10.1101/108944

**Authors:** Dobromir Rahnev

## Abstract

How are perceptual decisions made? The answer to this seemingly simple question necessitates that we specify the nature of perceptual representations on which decisions are based. Recent work has taken for granted that the representation at the decision stage consists of a full probability distribution over all possible stimuli. However, to date, no empirical evidence has supported this assumption. Here I present five possible perceptual representation schemes that allow the extraction of different levels of sensory uncertainty. I review the empirical evidence from both continuous and discrete judgments and show that, at present, only the most primitive scheme based on a single point estimate can be rejected. In other words, at least four different representational schemes are consistent with the available data and therefore full probability distributions cannot be assumed. There is an urgent need for empirical research to adjudicate between these theoretical possibilities.

## Introduction

Sometimes what we see is ambiguous or unclear. In such cases, we literally have to decide what is in front of our eyes. The idea that perception is a process of inference is as old as the study of perception itself^1^. Nevertheless, despite a lot of progress, it remains unknown what type of internal representation the observer forms in order to make a decision.

Significant amount of theoretical work has assumed that perceptual decisions are based on a representation of a full probability distribution over possible stimuli^2–19^. The idea was first developed about 20 years ago^13,14,20^ and has come to dominate our understanding of how the brain performs perceptual inference. Two particularly prominent theories include probabilistic population codes^7^ and neural sampling^10^ (**Box 1**).

Despite its popularity, the notion that full probability distributions are used to make perceptual decisions is not based on any direct evidence. Instead, proponents have often relied on theoretical arguments such as stating that the brain must represent optimal solutions and therefore full probability distributions are needed^12,18^. The empirical evidence cited in support of full probability distributions comes from just two classes of findings: cue combination is often optimal^22–25^, and priors and expectations bias perceptual decisions^26–32^.

This paper explains why none of the empirical evidence to date actually necessitates full probability distributions at the decision stage. It also critically examines the theoretical arguments for and against full probability distribution. To facilitate the discussion, I start by discussing possible schemes that support different levels of uncertainty representation.

#### Box1.Probabilistic population codes and neural sampling models

It has long been recognized that neurons in many sensory cortices have a specific tuning curve that controls how they respond to stimuli that lie on a continuum^21^.The collection of responses by many neurons forms a population response from which one can extract a full probability distribution over possible stimuli^13^.

At the same time, the representation of a given stimulus is different in different stages of the visual hierarchy. Therefore, the representation in sensory cortex may not be the same as the representation in decision-related areas. In other words, even if full distributions *could* be extracted from the population response in sensory areas such as V1 or MT, it does not follow that this information is necessarily available at the decision stage.

Two popular models propose that full probability distributions are in fact available on the decision stage. These models are probabilistic population codes^7^ and neural sampling with a large number of samples^10^. Probabilistic population codes (Panel A below) imply that the sensory population code can be used directly to form perceptual decisions. Neural sampling models (Panel B below), on the other hand, propose that neurons take discrete samples from the stimulus in small time intervals. These samples are then combined into a full distribution, provided that enough samples can be taken. Note that taking a single sample results in a point estimate (equivalent to Scheme 1 below), while taking only a few samples results in a very imperfect representation of the full distribution (conceptually similar, but not identical, to Schemes 3 and 4 below).

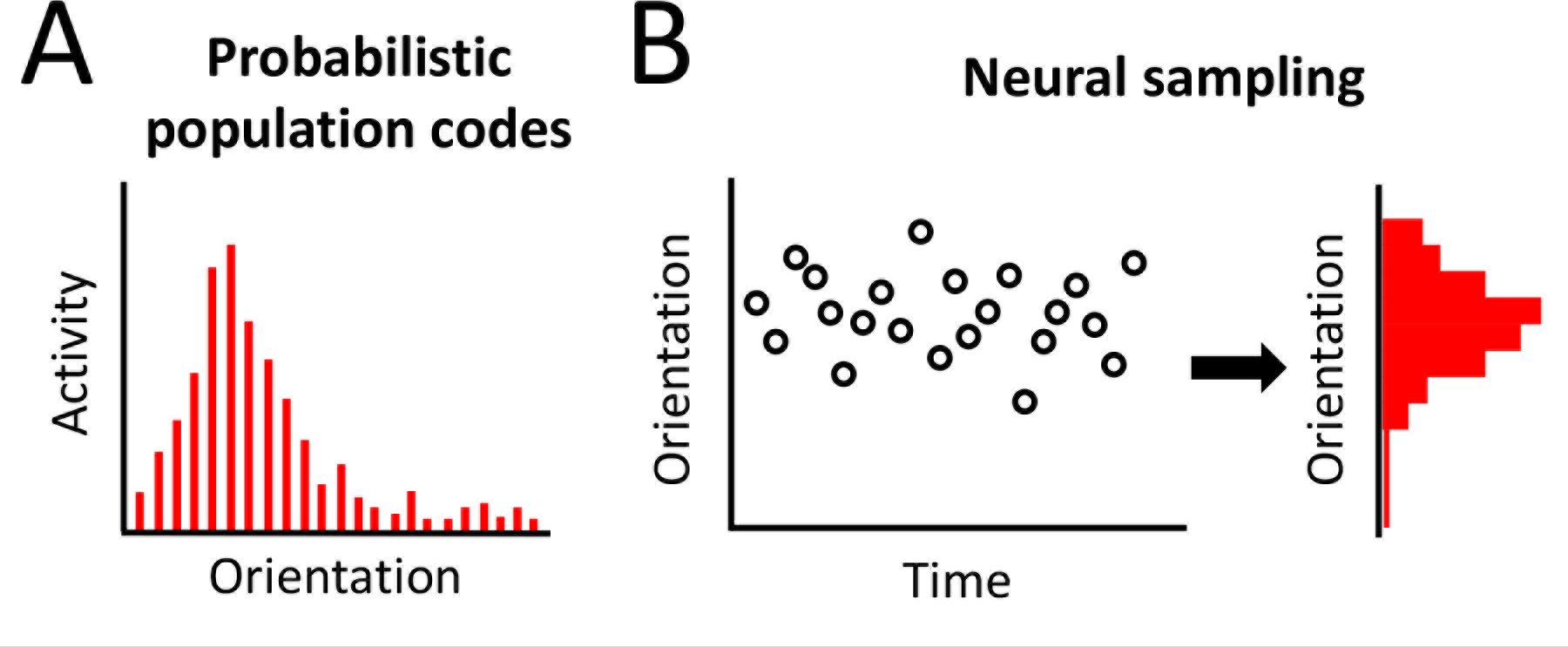

Two prominent theories postulate the existence of full probability distributions at the decision stage. (A) Probabilistic population codes assume that a population code is formed by examining the activity (within a certain time window) of neurons tuned to different values of a variable such as orientation. (B) Neural sampling models assume that at every time step, the brain extracts a single sample based on the whole neural population (left). Multiple samples are then joined together to form a distribution (right). Thus, through very different means, both probabilistic population codes and neural sampling postulate the existence of a full probability distribution (highlighted in red for each model).

### Five Schemes for the Nature of the Perceptual Representation

Many researchers appear to assume that perceptual representations consist either of simple point estimates or full probability distributions. In fact, there are a number of “intermediate” possibilities that often remain neglected. Here I present five different schemes for the nature of the perceptual representation at the decision stage. To compare between them, I consider separately continuous estimation and discrete choice tasks (**Box 2**).

As an example of a continuous estimation task, I will use the estimation of the direction of a single moving bar. The moving bar gives rise to a population code which can be represented as a sensory distribution in orientation space (**Figure 1A**). The question is what part of this population code is available for making the final decision.

As an example of a discrete choice task, I will use the decision about which of three possible stimuli - a face, a house, or a tool - was presented. These stimuli are known to be processed by relatively specialized areas within the ventral visual stream. The sensory evidence for each of these three possibilities can then be extracted from the level of activation of the corresponding brain areas (**Figure 2A**). Again, the question is what part of these activation levels are available for making the final decision.

##### Box 2. Continuous and discrete representations

In everyday life, we make decisions about both continuous quantities (e.g., how tall is this person, how fast is this car driving, how long has it been since lunch) and discrete choices (e.g., who is this person next to me in this old photo, what animal left these marks in the snow, is she pregnant). Theories regarding the nature of the perceptual representation at the decision stage have typically focused exclusively on tasks that require the estimation of continuous quantities. Part of the reason for this limitation is that the literature on discrete decisions is dominated by 2-choice tasks where different representational schemes are hard to distinguish. Nevertheless, truly understanding the nature of the perceptual representations necessitates that we specify the representation for both continuous and discrete quantities. The five schemes presented here are each applied to both types of judgments. It is possible that the brain representations for continuous and discrete judgments are qualitatively different, but, without any evidence for such a possibility, this paper assumes that each scheme should apply to both types of decisions.

#### Scheme 1. Single Point Estimate

The first possibility is that the perceptual representation at the decision stage consists of a single point estimate. In the continuous estimation task, the direction of a moving bar would be represented as a single orientation (e.g., 70°; **Figure 1B**). This point estimate could be based on the mean, median, mode, or any other statistic derived from the sensory distribution. In the discrete choice task, the representation could simply consist of the most likely stimulus (e.g., face; **Figure 2B**). The critical point is that in both cases only a single point estimate is passed onto the decision stage.

By assuming the presence of a single point estimate, Scheme 1 does not allow the estimation of sensory uncertainty. Indeed, on each trial, observers only have access to a single value with no way of extracting any information about how likely or unlikely that value is.

#### Scheme 2. Point Estimates for Task and Related Quantities

The second possibility is that the perceptual representation at the decision stage consists of multiple point estimates. For example, in addition to the point estimate from Scheme 1, people may additionally estimate other potentially relevant quantities such their own decision time and attentional state (**Figures 1C**, **2C**).

Scheme 2 allows for an indirect estimate of the true level of uncertainty. For example, decisions made more quickly and with higher attention are more likely to be correct and therefore would be associated with lower uncertainty. Thus, in Scheme 2, uncertainty about the stimulus does not stem from the sensory representation of the stimulus but of other, potentially related quantities.

#### Scheme 3. Point Estimate with Strength of Evidence

The third possibility is that the perceptual representation at the decision stage consists of a point estimate complemented by the strength of evidence for the point estimate. The strength of evidence here comes from the amount of evidence for the chosen option independent of all other information. For example, in the continuous estimation task, the direction of a moving bar could be represented by a point estimate (e.g., 70°), together with the level of activity of the neuron representing this point estimate (**Figure 1D**). Similarly, in the discrete choice task, the representation at the decision stage may consist of the identity of the stimulus with highest activity (e.g., face), together with the level of that activity in face-selective regions (**Figure 2D**).

Unlike in Scheme 2, the additional information (e.g., the strength of evidence) in Scheme 3 stems directly from the sensory distribution. However, because only partial information is considered (the activity level for the winning choice but not for the other choices), the uncertainty representation is incomplete.

#### Scheme 4. Partial Distribution

The fourth possibility is that the perceptual representation at the decision stage consists of a point estimate, as well as at least one higher moment of the sensory distribution (e.g., variance, skewness, kurtosis, etc.). For simplicity, in continuous estimation tasks, I will equate Scheme 4 with representing the mean and standard deviation. On a single trial, the direction of a moving bar may therefore be represented as 70° ± 30° (**Figure 1E**). Discrete tasks are more complicated since the options (e.g., face, house, tool) do not have an intrinsic relationship like the different orientations in a motion estimation task. In such discrete tasks, Scheme 4 can be seen as representing a point estimate, together with the entropy H(*X*) of the sensory representation (**Figure 2E**).

Scheme 4 features partial estimation of sensory uncertainty. This scheme carries more information than Schemes 1-3 since, unlike them, it takes into account the whole sensory distribution.

#### Scheme 5. Full Distribution

The final possibility is that the perceptual representation at the decision stage consists of a full probability distribution. In other words, the full sensory distribution is either replicated or simply accessed at the decision stage (**Figures 1F**, **2F**). Unlike Schemes 1-4, no point estimate is given a privileged representation; rather, all information is made available for further computations.

This scheme is the only one that allows the complete estimation of sensory uncertainty. The difference with Schemes 1-4 is particularly great in cases of discrete representations, as well as in complex continuous cases such as skewed or bimodal continuous distributions. Scheme 5 is also the only scheme that allows for fully optimal decisions on every trial. As described above, these features have made this scheme very popular among computational neuroscientists^2–19^.

**Figure 1.**
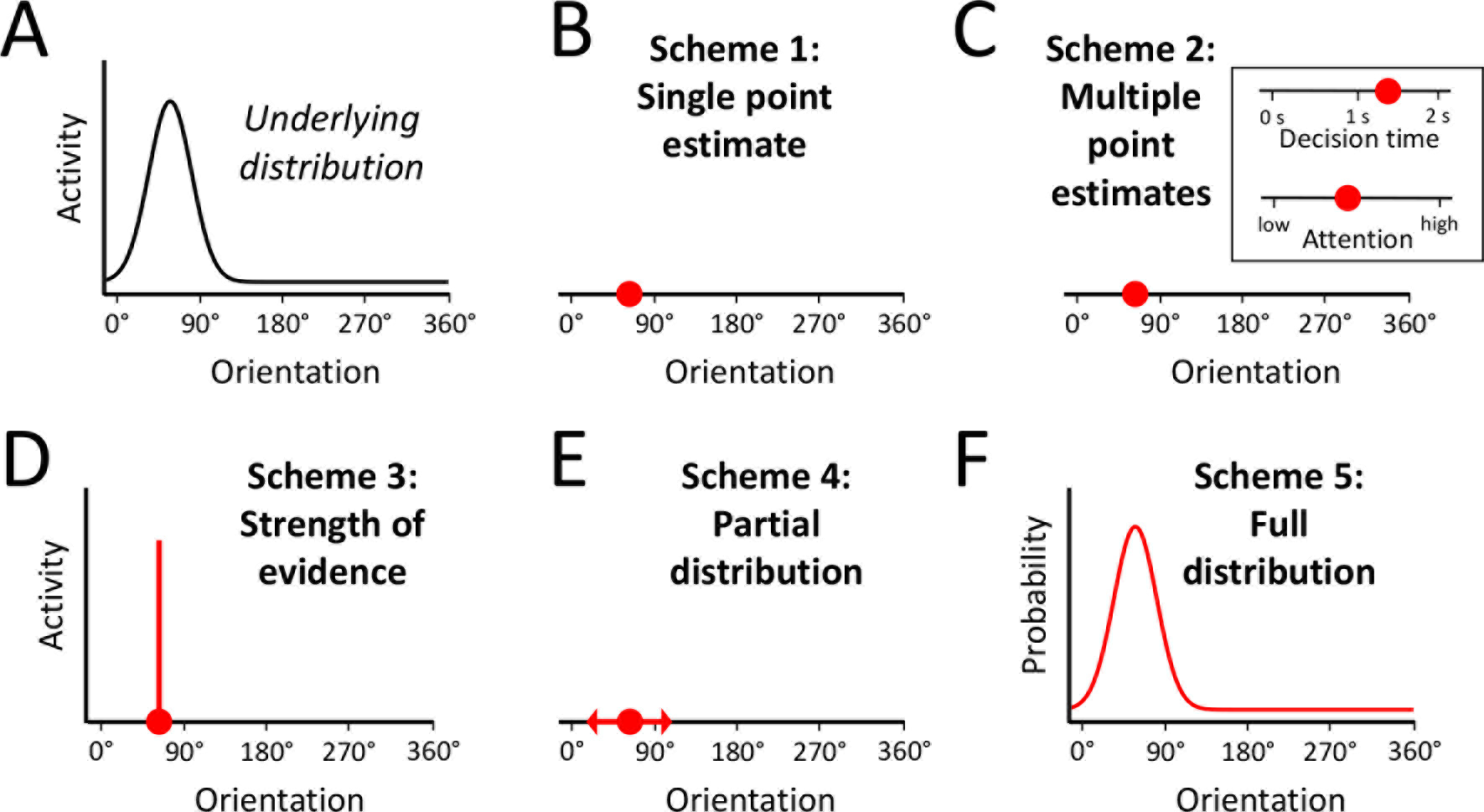
Five schemes for the nature of the perceptual representation in continuous tasks. (A) A moving bar creates a Gaussian distribution in motion-sensitive MT neurons tuned to different orientations. (B) Scheme 1: Single point estimate. Only a single point estimate of the direction of motion of the bar is extracted at the decision stage. (C) Scheme 2: Multiple point estimates. Several point estimates are extracted for variables relevant to the task (e.g., the inset shows estimates of decision time and attentional state). (D) Scheme 3: Strength of evidence. A point estimate and a strength-of-evidence value are extracted. The strength of evidence corresponds to the height of the distribution at the point estimate. (E) Scheme 4: Partial probability distribution. The mean and standard deviation of the distribution of neuronal activity are extracted. (F) Scheme 5: Full probability distribution. The whole distribution of neuron activity is used but is transformed from “activity” to “probability” space. The information extracted in each scheme is highlighted in red.

**Figure 2.**
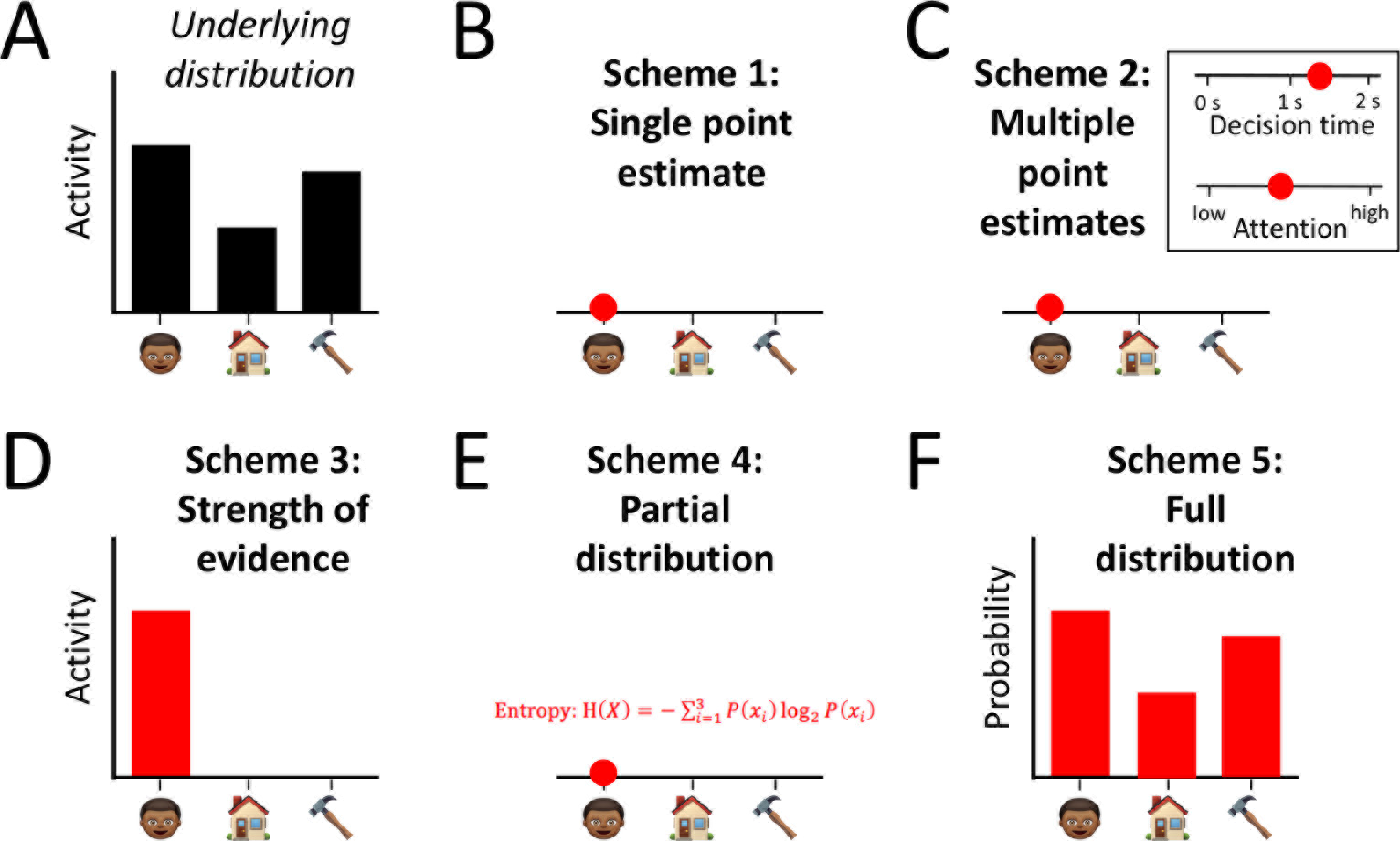
Five schemes for the nature of the perceptual representation in discrete tasks. (A) An ambiguous stimulus creates various amount of activity in sensory cortex coding for faces, houses, and tools. (B) Scheme 1: Single point estimate. Only a single guess about the identity of the ambiguous stimulus is extracted at the decision stage. (C) Scheme 2: Multiple point estimates. Several point estimates are extracted for variables relevant to the task (e.g., the inset shows estimates of decision time and attentional state). (D) Scheme 3: Strength of evidence. A single guess together with the activity in sensory cortex corresponding to that guess are extracted. (E) Scheme 4: Partial probability distribution. A single guess together with the entropy associated with it are extracted. (F) Scheme 5: Full probability distribution. The complete sensory distribution is used but is transformed from “activity” to “probability” space. The information extracted in each scheme is highlighted in red.

### Empirical Evidence

Having presented five possible schemes of the perceptual representation at the decision level, I now explore the empirical evidence typically invoked in support of Scheme 5. Remarkably, all arguments are indirect and at least 15 years old. Even though not typically considered in this context, I also examine the evidence from confidence ratings. In each case, I show that rather than providing support for Scheme 5, the empirical evidence is consistent with all of Schemes 2-5, and even place strong constraints on Scheme 5.

#### Cue Combination

Cue combination occurs when two or more pieces of information are combined to form a single decision^33^. Such tasks are typically performed with stimuli that permit continuous estimation. For example, the length of a bar could be estimated based on a combination of visual and haptic information^23^. When the information from each of these sensory modalities is noisy, their evidence is combined in order to arrive at a better estimate than either sense can afford by itself.

Cue combination studies often find near optimal integration^22–25^. Such findings have been cited as providing the strongest support for the existence of a full probability distribution as in Scheme 5^2–11^. Indeed, the existence of a full probability distribution easily explains optimal cue combination^7^. Importantly, Scheme 4 also naturally fits with optimal cue combination since such combination only requires the representation of the distributions’ mean and standard deviation. In fact, assuming Gaussian variability of the underlying sensory distribution, Scheme 4 becomes equivalent to Scheme 5 when applied to cue combination.

What is less appreciated is that Scheme 3, and perhaps even by Scheme 2, can also explain near optimal performance in cue combination studies. Scheme 3 requires subjects to weight each stimulus’ point estimate by the strength-of-evidence value associated with it. In cases of Gaussian variability, the height of the distribution is directly related to its standard deviation (for a normalized distribution), so Scheme 3 can support optimal performance. Scheme 2 has more difficulties with near optimal cue combination because of its indirect representation of sensory uncertainty. Still, occasional near optimal performance is possible when parameters such as decision time or attentional state are strongly correlated with performance. Thus, only Scheme 1 is completely inconsistent with near optimal cue combination.

On the other hand, discussions of cue combination studies often ignore the fact that many such studies find substantial suboptimalities^34–42^ (reviewed in ^43^). These studies typically report that one of the cues was weighted more than its reliability, relative to the other cue. Such findings are surprising if indeed perceptual decisions are based on full probability distributions (Scheme 5). On the other hand, it is more natural to expect that Schemes 2 and 3 (and, to a lesser degree, Scheme 4) will sometimes lead to suboptimal cue combination since they rely on heuristics.

#### Priors, Expectations, and Confidence

Subjects are able to seamlessly incorporate priors acquired by experience^26,27^ or provided by an experimenter^28–32^ to improve their perceptual decisions. Similarly, in the absence of any prior, subjects can provide confidence ratings that reflect the probability of being correct^44,45^. (Priors and expectations are traditionally studied in the context of both continuous and discrete tasks, while confidence studies typically focus on discrete tasks.) These abilities constitute strong evidence against Scheme 1. Indeed, this scheme does not feature any information on which appropriate use of priors or meaningful confidence judgments could be based. Schemes 2-5 easily explain how priors can be incorporated into decisions or meaningful confidence ratings can occur: the influence of priors should decrease and confidence should increase with higher attentional (Scheme 2), higher strength of evidence (Scheme 3),lower standard deviations (Scheme 4), or narrower distributions (Scheme 5).

On the other hand, a number of suboptimalities exist in both the use of priors and expectations, as well as in confidence ratings^43^. For example, subjects typically underuse priors that are explicitly provided by the experimenter^28–32^. They are also frequently either under-or over-confident and are rarely able to calibrate their confidence properly^44^,^46–52^. Even more importantly, a large number of studies have reported conditions matched on accuracy that produce different levels of confidence^44^,^53–66^. These findings strongly suggest the existence of heuristics and biases in both the use of priors and in confidence computation.

Similar to cue combination tasks, the suboptimalities in combining priors with sensory information and in confidence ratings present a problem for Scheme 5. Indeed, at least in principle, the presence of complete probability distributions should allow for fully optimal computations. On the other hand, Schemes 2-4 fit much more naturally with these findings of suboptimality since the probability of being correct has a complex relationship with their measures of uncertainty. These considerations explain why Schemes 2-4 would predict the existence of biases in the use of priors and confidence ratings.

### Theoretical Arguments

As noted in the beginning, proponents of full probability distributions at the decision stage often motivate their stance with theoretical arguments rather than empirical evidence. Two main arguments are explicitly or implicitly advanced.

First, computational neuroscientists often prefer to build normative models of how the visual system should or could deal with uncertainty. Full distributions are best for normative computations. The belief that normative solutions are necessary is further fueled by a number of findings of close to optimal performance^22–25^. However, many other theorists have argued that there is no a priori reason to expect that the brain implements normative solutions, especially in complex situations^67–69^. Further, an extensive literature shows that suboptimal behavior has been observed in virtually every perceptual task that lends itself to optimality analysis^43^. On the balance, it is hard to argue that a mixture of optimal and suboptimal performance points to representations that are specifically designed for optimality (like Scheme 5). One can argue that in many situations the information from the full distributions is not used optimally. While this is certainly possible, this argument begs the question as to why full distributions would be represented in the first place if they are often not used properly.

Second, proponents of full distributions often point out that a single point estimate cannot explain many behavioral effects^2,3,5,10,19^, thus rejecting Scheme 1. Scheme 5 is then presented as the only possible alternative. In this view, any finding that the brain can extract uncertainty estimates from sensory representations is taken as evidence that decisions are “Bayesian” or “probabilistic” (**Box 3**), which is later equated with full probability distributions. As demonstrated by Schemes 2-4, many other alternatives exist.

Not only do theoretical arguments for Scheme 5 appear lacking, but there are at least two arguments *against* the existence of full probability distributions. First, real-life perception comes with an explosion in computational complexity. Such complexity virtually guarantees that decisions will be based on heuristics rather than fully principled computations^67–69^. Even supporters of full probability distributions admit that complex situations call for simplified computations^6^. Perception evolved to serve us in real life rather than in the laboratory. Thus, complex conditions are the norm rather than the exception for the brain. If heuristics are necessary anyway, then perceptual representations that allow precise estimation of uncertainty but are mostly applicable in very simple situations may be an unnecessary luxury.

Second, perceptual judgments are known to become more optimal with practice^44,70,71^. Such findings fit well with Schemes 2 and 3. The reason is that in both of these schemes, the additional quantities (beyond the point estimate) such as decision time and strength of evidence may predict one’s accuracy differently for different tasks. Thus, both Schemes 2 and 3 require learning to calibrate how these additional quantities should be used. However, it is less clear how and why learning would make judgments more optimal in Schemes 4 and 5.

##### Box 3. Are perceptual decisions Bayesian? Are they probabilistic?

The possibility that perceptual representations at the decision stage do not consist of full probability distributions has relevance to theories about “Bayesian brains,” “Bayesian computation,” “probabilistic approach,” “probabilistic computation,” and “probabilistic brains”^2–9^. These concepts sound similar but are often used with very different meanings.

Are perceptual decisions Bayesian? Decisions are Bayesian as long as they follow Bayes’ theorem. Much evidence suggests that they do, although specific computations may deviate from optimality for a number of reasons^43^. Importantly, many Bayesian models only require a single point estimate and build the Bayesian machinery around inferring how the point estimate varies over trials. Thus, all five schemes are fully consistent with the notion that perceptual decisions are Bayesian. In some cases, however, the term “Bayesian” is equated with “optimal.” As pointed out above, it does not appear that all perceptual decisions are optimal, so under this more restrictive definition, perceptual decisions should not be generally classified as Bayesian.

Are perceptual decisions probabilistic? The term “probabilistic” is even more challenging. Ma^2^ distinguishes between *probabilistic models* (in which trial-to-trial observations are stochastic; such models can feature representations consistent with all five schemes) and *models of probabilistic computation* (which require the representation of at least two moments of the sensory distribution; such models are only consistent with Schemes 4 and 5). Nevertheless, Ma’s terminology is not widely used. Other papers equate phrases such as the “probabilistic approach”^6^ and representing stimuli in a “probabilistic manner”^10^ with the existence of the full probability distributions from Scheme 5. Thus, rejecting Scheme 5 means rejecting the notion of probabilistic decisions in some but not other definitions of the term.

So,are perceptual decisions Bayesian and/or probabilistic? It depends on what one means when using these terms. There is strong evidence for stochasticity and use of Bayes’ theorem in perception but no evidence for full probability distributions or complete optimality. Due to the ambiguity in existing terminology, future work could refer to Schemes 1-5 in order to clarify the exact concept researchers seek to advance. Increased clarity may also help avoid common logical traps such as stating that the perceptual decisions follow Bayes’ theorem (true) and concluding that full probability distributions are necessarily needed (false).

### What Empirical Evidence Can Resolve the Issue?

It is safe to say that Scheme 1 has been thoroughly debunked. However, it is much harder to distinguish the rest of the schemes. The critical question is whether the perceptual representation features *anything* more than a point estimate with some uncertainty estimate (as in Schemes 2-4). Scheme 5 can of course come in different flavors with various amounts of precision in the representation. Nevertheless, there are at least two ways of adjudicating between Scheme 5 (even if instantiated with relatively low precision) and Schemes 2-4.

First, one can employ stimuli that produce continuous but non-Gaussian distributions of sensory evidence. As pointed out above, especially useful would be bi-or tri-modal distributions. For such distributions, Schemes 2-4 can only represent a single peak in the distribution, while Scheme 5 can represent all peaks simultaneously.

Second, one can employ discrete choice tasks with multiple alternatives. Again, Schemes 2-4 can represent only the most likely option (with an added uncertainty estimate), while only Scheme 5 can represent the evidence for all options. Note that more than two alternatives would be needed if Scheme 5 is to make qualitatively different predictions than Schemes 2-4.

Both options above are constructed such that Scheme 5 represents meaningfully more information than Schemes 2-4. Clever experimental designs can then be leveraged in order for Schemes 2-4 vs. Scheme 5 to make different predictions. What is critical to appreciate is that Scheme 5 can and should be empirically adjudicated from Schemes 2-4.

## Conclusion

The nature of the perceptual representation at the decision level is still a mystery. Many theorists have strongly favored the idea that decisions are based on full probability distributions over possible choices. However, closer scrutiny demonstrates that, at present, there is no evidence for such a proposition. Instead, the empirical data strongly rejects only representations based on a single point estimate (Scheme 1). At least three other possible representations based on a point estimate and various types of additional information (Scheme 2-4) cannot currently be clearly distinguished from full probability distributions (Scheme 5).

Is it possible that Scheme 5 is the correct one after all? Absolutely. However, before empirical evidence is actually leveraged in support of it (thus falsifying Schemes 2
, we cannot and should not simply assume the existence of full probability distributions at the decision level. There is an urgent need to address this question empirically before more theoretical work on the subject is done.

## Acknowledgments

I am thankful to Luigi Acerbi, David Carmel, Michael Landy, Wei Ji Ma, and Annelinde Vandenbroucke for helpful comments. This work was funded by a startup grant from the Georgia Institute of Technology.

## References

1. Helmholtz, H. L. F. Treatise on physiological optics. (Thoemmes Continuum, 1856).

2. Ma, W. J. Signal detection theory, uncertainty, and Poisson-like population codes. Vision Res. 50, 2308–2319 (2010).

3. Ma, W. J. & Jazayeri, M. Neural Coding of Uncertainty and Probability. Annu. Rev. NeuroSci. 37, 205–220 (2014).

4. Ma, W. J. Organizing probabilistic models of perception. Trends Cogn. Sci. 16, 511–8 (2012).

5. Knill, D. C. & Pouget, A. The Bayesian brain: the role of uncertainty in neural coding and computation. Trends NeuroSci. 27, 712–9 (2004).

6. Pouget, A., Beck, J. M., Ma, W. J. & Latham, P. E. Probabilistic brains: knowns and unknowns. Nat. NeuroSci. 16, 1170–8 (2013).

7. Ma, W. J., Beck, J. M., Latham, P. E. & Pouget, A. Bayesian inference with probabilistic population codes. Nat. NeuroSci. 9, 1432–8 (2006).

8. Drugowitsch, J. & Pouget, A. Probabilistic vs. non-probabilistic approaches to the neurobiology of perceptual decision-making. Curr. Opin. Neurobiol. 22, 963–9 (2012).

9. Beck, J. M., Ma, W. J., Pitkow, X., Latham, P. E. & Pouget, A. Not noisy, just wrong: the role of suboptimal inference in behavioral variability. Neuron 74, 30–9 (2012).

10. Fiser, J., Berkes, P., Orbán, G. & Lengyel, M. Statistically optimal perception and learning: from behavior to neural representations. Trends Cogn. Sci. 14, 119–30 (2010).

11. Berkes, P., Orbán, G., Lengyel, M. & Fiser, J. Spontaneous cortical activity reveals hallmarks of an optimal internal model of the environment. Science (80-.). 331, 83–7 (2011).

12. Jazayeri, M. & Movshon, J. A. Optimal representation of sensory information by neural populations. Nat. NeuroSci. 9, 690–6 (2006).

13. Zemel, R. S., Dayan, P. & Pouget, A. Probabilistic Interpretation of Population Codes. Neural Comput. 10, 403–430 (1998).

14. Pouget, A., Dayan, P. & Zemel, R. Information processing with population codes. Nat. Rev. NeuroSci. 1, 125–132 (2000).

15. Sahani, M. & Dayan, P. Doubly distributional population codes: Simultaneous representation of uncertainty and multiplicity. Neural Comput. 15, 2255–2279 (2003).

16. Haefner, R. M., Berkes, P. & Fiser, J. Perceptual Decision-Making as Probabilistic Inference by Neural Sampling. Neuron 90, 649–660 (2016).

17. Sanborn, A. N. & Chater, N. Bayesian Brains without Probabilities. Trends Cogn. Sci. 20, 883–893 (2016).

18. Beck, J., Pouget, A. & Heller, K. A. Complex Inference in Neural Circuits with Probabilistic Population Codes and Topic Models. in Advances in Neural Information Processing Systems 25 3059–3067 (2012). at <https://papers.nips.cc/paper/4555-complex-inference-in-neural-circuits-with-probabilistic-population-codes-and-topic-models>

19. Pouget, A., Dayan, P. & Zemel, R. S. Inference and Computation with Population Codes. Annu. Rev. NeuroSci. 26, 381–410 (2003).

20. Zemel, R. S. & Dayan, P. Distributional Population Codes and Multiple Motion Models. in Advances in Neural Information Processing Systems 11 174–180 (MIT Press, 1999).

21. Adrian, E. D. The impulses produced by sensory nerve endings. J. Physiol. 61, 49–72 (1926).

22. Alais, D. & Burr, D. The ventriloquist effect results from near-optimal bimodal integration. Curr. Biol. 14, 257–62 (2004).

23. Ernst, M. O. & Banks, M. S. Humans integrate visual and haptic information in a statistically optimal fashion. Nature 415, 429–33 (2002).

24. Gu, Y., Angelaki, D. E. & DeAngelis, G. C. Neural correlates of multisensory cue integration in macaque MSTd. Nat. NeuroSci. 11, 1201–1210 (2008).

25. van Beers, R. J., Sittig, A. C. & Denier van der Gon, J. J. How humans combine simultaneous proprioceptive and visual position information. Exp. brain Res. 111, 253–61 (1996).

26. Weiss, Y., Simoncelli, E. P. & Adelson, E. H. Motion illusions as optimal percepts. Nat. NeuroSci. 5, 598–604 (2002).

27. Stocker, A. A. & Simoncelli, E. P. Noise characteristics and prior expectations in human visual speed perception. Nat. NeuroSci. 9, 578–85 (2006).

28. Ackermann, J. F. & Landy, M. S. Suboptimal decision criteria are predicted by subjectively weighted probabilities and rewards. Atten. Percept. Psychophys. 77, 638–658 (2015).

29. Rahnev, D., Lau, H. & De Lange, F. P. Prior expectation modulates the interaction between sensory and prefrontal regions in the human brain. J. NeuroSci. 31, 10741–10748 (2011).

30. de Lange, F. P., Rahnev, D., Donner, T. H. & Lau, H. Prestimulus Oscillatory Activity over Motor Cortex Reflects Perceptual Expectations. J. NeuroSci. 33, 1400–1410 (2013).

31. Summerfield, C. & Koechlin, E. Economic value biases uncertain perceptual choices in the parietal and prefrontal cortices. Front. Hum. NeuroSci. 4, 208 (2010).

32. Ulehla, Z. J. Optimality of perceptual decision criteria. J. Exp. Psychol. 71, 564–569 (1966).

33. Sensory Cue Integration. (Oxford University Press, 2011).

34. Battaglia, P. W., Jacobs, R. A. & Aslin, R. N. Bayesian integration of visual and auditory signals for spatial localization. J. Opt. Soc. Am. A, Opt. image Sci. 20, 1391–1397 (2003).

35. Maiworm, M. & Röder, B. Suboptimal Auditory Dominance in Audiovisual Integration of Temporal Cues. Tsinghua Sci. Technol. 16, 121–132 (2011).

36. Burr, D., Banks, M. S. & Morrone, M. C. Auditory dominance over vision in the perception of interval duration. Exp. Brain Res. 198, 49–57 (2009).

37. Fetsch, C. R., Pouget, A., Deangelis, G. C. & Angelaki, D. E. Neural correlates of reliability-based cue weighting during multisensory integration. Nat. NeuroSci. 15, 146–154 (2012).

38. Prsa, M., Gale, S. & Blanke, O. Self-motion leads to mandatory cue fusion across sensory modalities. J. Neurophysiol. 108, 2282–2291 (2012).

39. Battaglia, P. W., Kersten, D. & Schrater, P. R. How haptic size sensations improve distance perception. PLoS Comput. Biol. 7, e1002080 (2011).

40. Rosas, P., Wagemans, J., Ernst, M. O. & Wichmann, F. A. Texture and haptic cues in slant discrimination: reliability-based cue weighting without statistically optimal cue combination. J. Opt. Soc. Am. A, Opt. image Sci. 22, 801–809 (2005).

41. Knill, D. C. & Saunders, J. A. Do humans optimally integrate stereo and texture information for judgments of surface slant? Vision Res. 43, 2539–2558 (2003).

42. Rosas, P., Wichmann, F. A. & Wagemans, J. Texture and object motion in slant discrimination: failure of reliability-based weighting of cues may be evidence for strong fusion. J. Vis. 7, 3 (2007).

43. Rahnev, D. & Denison, R. Suboptimality in perception. bioRxiv (2016). at “http://biorxiv.org/content/early/2016/06/22/060194.abstract”

44. Baranski, J. V & Petrusic, W. M. The calibration and resolution of confidence in perceptual judgments. Percept. Psychophys. 55, 412–28 (1994).

45. Fleming, S. M. & Lau, H. How to measure metacognition. Front. Hum. NeuroSci. 8, (2014).

46. Adams, J. K. A confidence scale defined in terms of expected percentages. Am. J. Psychol. 70, 432–6 (1957).

47. Dawes, R. M. in Similarity and choice: Papers in honor of Clyde Coombs (eds. Lantermann, E. D. & Feger, H.) 327–345 (Han Huber, 1980).

48. Harvey, N. Confidence in judgment. Trends Cogn. Sci. 1, 78–82 (1997).

49. Keren, G. On the ability of monitoring non-veridical perceptions and uncertain knowledge: Some calibration studies. Acta Psychol. (Amst). 67, 95–119 (1988).

50. Koriat, A. Subjective confidence in perceptual judgments: a test of the selfconsistency model. J. Exp. Psychol. Gen. 140, 117–139 (2011).

51. Björkman, M., Juslin, P. & Winman, A. Realism of confidence in sensory discrimination: the underconfidence phenomenon. Percept. Psychophys. 54, 75–81 (1993).

52. Winman, A. & Juslin, P. Calibration of sensory and cognitive judgments: Two different accounts. Scand. J. Psychol. 34, 135–148 (1993).

53. Vickers, D. & Packer, J. Effects of alternating set for speed or accuracy on response time, accuracy and confidence in a unidimensional discrimination task. Acta Psychol. (Amst). 50, 179–97 (1982).

54. Kiani, R., Corthell, L. & Shadlen, M. N. Choice Certainty Is Informed by Both Evidence and Decision Time. Neuron 84, 1329–1342 (2014).

55. Rahnev, D. et al. Attention induces conservative subjective biases in visual perception. Nat. NeuroSci. 14, 1513–1515 (2011).

56. Rahnev, D., Bahdo, L., de Lange, F. P. & Lau, H. Prestimulus hemodynamic activity in dorsal attention network is negatively associated with decision confidence in visual perception. J. Neurophysiol. 108, 1529–36 (2012).

57. Wilimzig, C., Tsuchiya, N., Fahle, M., Einhäuser, W. & Koch, C. Spatial attention increases performance but not subjective confidence in a discrimination task. J. Vis. 8, 1–10 (2008).

58. de Gardelle, V. & Mamassian, P. Weighting Mean and Variability during Confidence Judgments. PLoS One 10, e0120870 (2015).

59. Koizumi, A., Maniscalco, B. & Lau, H. Does perceptual confidence facilitate cognitive control? Atten. Percept. Psychophys. (2015). doi:10.3758/s13414-015-0843-3

60. Song, A., Koizumi, A. & Lau, H. in Behavioral Methods in Consciousness Research (ed. Overgaard, M.) (Oxford University Press, 2015).

61. Spence, M. L., Dux, P. E. & Arnold, D. H. Computations Underlying Confidence in Visual Perception. J. Exp. Psychol. Hum. Percept. Perform. 42, 671–682 (2016).

62. Zylberberg, A., Roelfsema, P. R. & Sigman, M. Variance misperception explains illusions of confidence in simple perceptual decisions. Conscious. Cogn. 27, 246–253 (2014).

63. Samaha, J., Barrett, J. J., Sheldon, A. D., LaRocque, J. J. & Postle, B. R. Dissociating Perceptual Confidence from Discrimination Accuracy Reveals No Influence of Metacognitive Awareness on Working Memory. Front. Psychol. 7, 851 (2016).

64. Vlassova, A., Donkin, C. & Pearson, J. Unconscious information changes decision accuracy but not confidence. Proc. Natl. Acad. Sci. 111, 16214–16218 (2014).

65. Navajas, J., Sigman, M. & Kamienkowski, J. E. Dynamics of visibility, confidence, and choice during eye movements. J. Exp. Psychol. Hum. Percept. Perform. 40, 1213–1227 (2014).

66. Rahnev, D., Koizumi, A., McCurdy, L. Y., D’Esposito, M. & Lau, H. Confidence Leak in Perceptual Decision Making. Psychol. Sci. 26, 1664–1680 (2015).

67. Gigerenzer, G. & Brighton, H. Homo Heuristicus: Why Biased Minds Make Better Inferences. Top. Cogn. Sci. 1, 107–143 (2009).

68. Juslin, P., Nilsson, H. & Winman, A. Probability theory, not the very guide of life. Psychol. Rev. 116, 856–874 (2009).

69. Simon, H. A. Rational choice and the structure of the environment. Psychol. Rev. 63, 129–138 (1956).

70. Maddox, W. T. & Bohil, C. J. Optimal classifier feedback improves cost-benefit but not base-rate decision criterion learning in perceptual categorization. Mem. Cognit. 33, 303–19 (2005).

71. Balci, F. et al. Acquisition of decision making criteria: reward rate ultimately beats accuracy. Atten. Percept. Psychophys. 73, 640–57 (2011).

